# Peptide-based drug predictions for cancer therapy using deep learning

**DOI:** 10.1101/2022.02.01.478580

**Authors:** Yih-Yun Sun, Tzu-Tang Lin, Wei-Chih Cheng, I-Hsuan Lu, Shu-Hwa Chen, Chung-Yen Lin

## Abstract

**Background:** Therapeutic drugs used in cancer treatment have ineffectiveness and resistance to drug action problems. Anticancer peptides (ACPs) are selective and toxic to cancer cells and quickly produced. Thus, ACPs can be a satisfactory substitute for therapeutic drugs. We developed AI4ACP, a user-friendly web-server ACP predictor that can predict the anticancer property of query peptides, thus promoting the discovery of peptides with anticancer activity.

**Result:** Our results revealed that the performance of our ACP predictor trained using the new ACP collection was superior to that of the available high-performance ACP predictors.

**Conclusions:** AI4ACP is a user-friendly web-server ACP predictor that can be used to determine whether a query sequence is an ACP. This tool can be beneficial for drug development for cancer treatment. AI4ACP is freely accessible at https://axp.iis.sinica.edu.tw/AI4ACP/

## Background

Therapeutic drugs currently used in cancer treatment have ineffectiveness and resistance to drug action, thus increasing side effects [1]. Cell membrane properties differ between cancer and healthy cells. The membrane fluidity of cancer cells is higher than that of healthy cells [2]; this protects cancer cells from cell lysis. In addition, cancer cells are characterized by a negatively charged surface [3]. Anticancer peptides (ACPs), a subset of antimicrobial peptides (AMPs), are selective and toxic to cancer cells because of their physicochemical properties and secondary structures. ACPs can be divided into two types based on their anticancer mechanism: molecular-targeting peptides and cancer-targeting peptides. Compared with therapeutic drugs, ACPs have higher specificity and selectivity and can easily bind to various targeting drugs. ACPs can be easily synthesized and produced and can thus serve as a new cancer treatment modality [1].

Some state-of-the-art predictors have been constructed using traditional machine learning methods such as support vector machine (SVM) for AntiCP [4] and iACP [5] and random forest (RF) for ACPred [6] and MLACP [7]. However, because traditional machine learning methods depend on manual feature extraction, their performance may be affected by the experience and knowledge of researchers.

Recently, deep learning models have been successfully applied in many fields (e.g., for the prediction of ACPs). A study, PTPD, used a combination of Word2Vec and a deep learning network (DNN) model [8]. With the availability of deep learning methods, researchers are currently not required to extract features manually. Instead, a machine extracts the features of data automatically. Moreover, with an increase in the number of ACPs confirmed recently, deep learning models have increased accurately.

To hasten the discovery of ACPs and improve the performance of ACP predictors, we built a deep learning model to detect peptides with anticancer activity. Our model was composed of peptide sequence encoding and machine learning. The protein-encoding stage involves using encoding methods, such as the analysis of amino acid and dipeptide composition, reported in previous studies. In this study, we used PC6 [9], a novel protein-encoding method, to convert a peptide sequence into a computational matrix, representing six physicochemical properties of each amino acid. We mainly applied the convolutional neural network in our model in the machine learning stage. Because of an increase in the number of ACPs confirmed recently, we could identify more ACP sequences and construct a highly accurate ACP prediction model.

## Implementation

### 1.1 Data collection and division

#### Positive data collection

We collected ACP sequences from four ACP and AMP databases: CancerPPD [10], DBAASP [11], DRAMP [12], and YADAMP [13]. In addition, we included sequences from the positive alternative set reported by Charoenkwan et al. 2021 [14]. We downloaded all peptides with anticancer activity from the four databases and previous studies. After excluding ACPs with unusual amino acids or a nonlinear structure, namely “B,” “Z,” “U,” “X,” “J,” “O,” “i,” and “-,” and duplicates between different databases, we obtained 2839 positive ACPs. Fig. 1 (Panel A) presents the length distribution of the 2839 ACPs; most of the sequences were shorter than 50 amino acids in length. Therefore, we excluded ACPs longer than 50 amino acids. Finally, 2815 ACP sequences were retained. Fig. 1 (Panel B) depicts the length distribution of the 2815 ACPs.

**Fig. 1.**
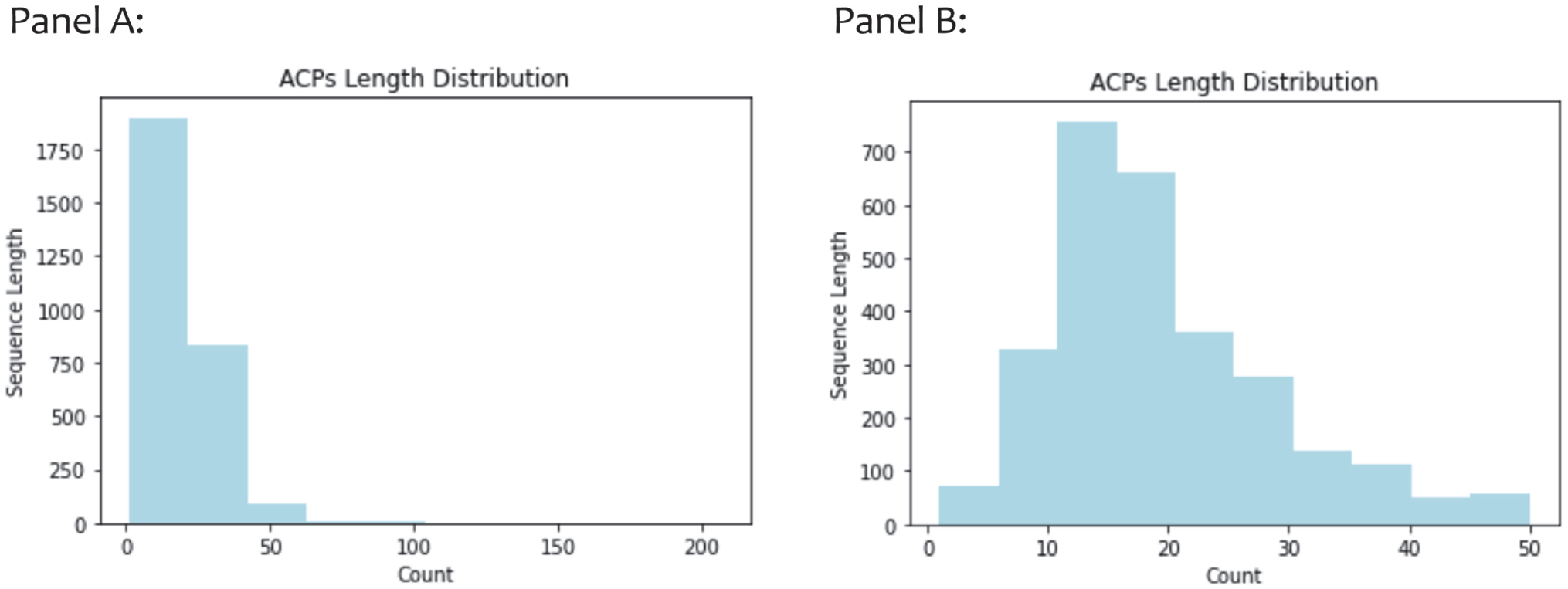
Histogram of the length of ACPs.

To ensure that the characteristics of the ACPs learned by the model were balanced, we filtered out the remaining ACPs sharing >99% sequence identity with existing ACPs by calculating the sequence identity by using CD-HIT [9]. A total of 2124 ACPs were included as positive data. To evaluate the performance of our model and compare it with that of other state-of-the-art predictors, we used 10% of all the positive data as the testing set after excluding sequences from the positive set of other predictors. Fig. 2 presents the detailed positive data collection and division process.

**Fig. 2.**
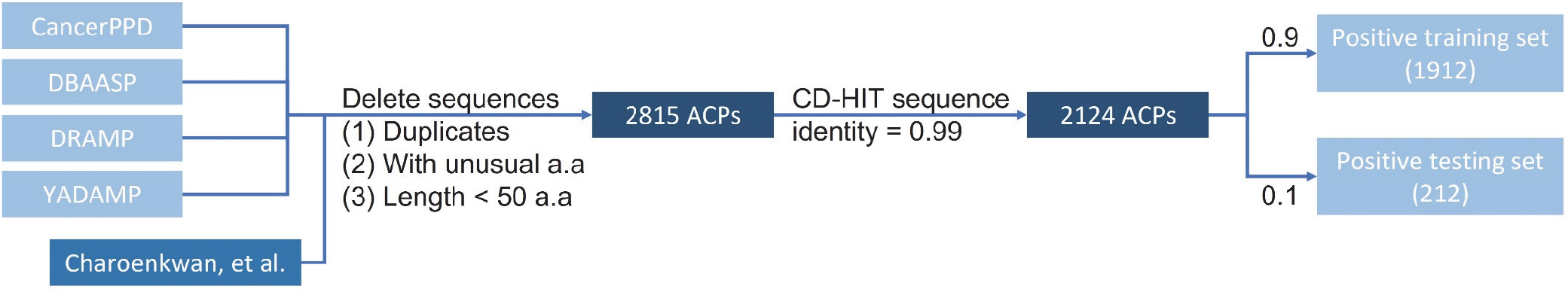
Positive data collection and division process.

#### Negative data collection

The negative data set consisted of 1062 non-ACP peptides from UniProt [15] and 1062 manually generated peptides. From UniProt, we collected peptides shorter than 50 amino acids in length and without anticancer, antiviral, antimicrobial, or antifungal activities. Manually generated peptides were randomly generated using the same length of the positive data set and 20 essential amino acids. Finally, we obtained 2124 sequences as the negative data set. We used 90% of the negative data set (1912 sequences) as the negative training set and the remaining 10% (212 sequences) as the negative testing set. Fig. 3 presents the detailed negative data collection and division process.

**Fig. 3.**
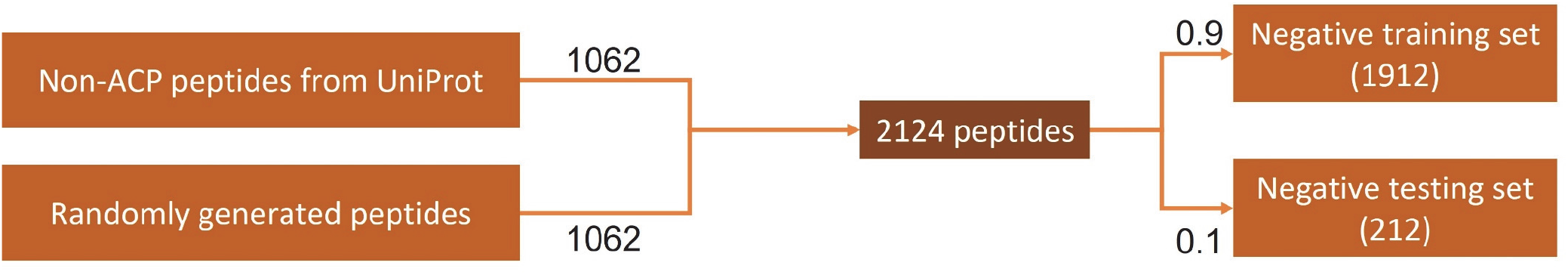
Negative data collection and division process.

### 1.2 Protein-encoding method

This study used the PC6 protein-encoding method [9] to convert a peptide sequence into a computational matrix. PC6 is a novel protein-encoding method that can encode a sequence based on both the order and physicochemical properties of the amino acids of the sequence. After benchmarking with other encoding methods, the PC6 encoding method exhibited the most satisfactory performance. Therefore, we applied PC6 in the encoding stage in our final prediction model.

### 1.3 Developing a deep learning model

We implemented Keras, a high-level API from Tensorflow, to construct and train a deep learning model. We first applied the PC6 protein-encoding method [9] to all sequences and converted them into 50 × 6 matrices. Fig. 4 presents the process of the PC6 protein-encoding method.

**Fig. 4.**
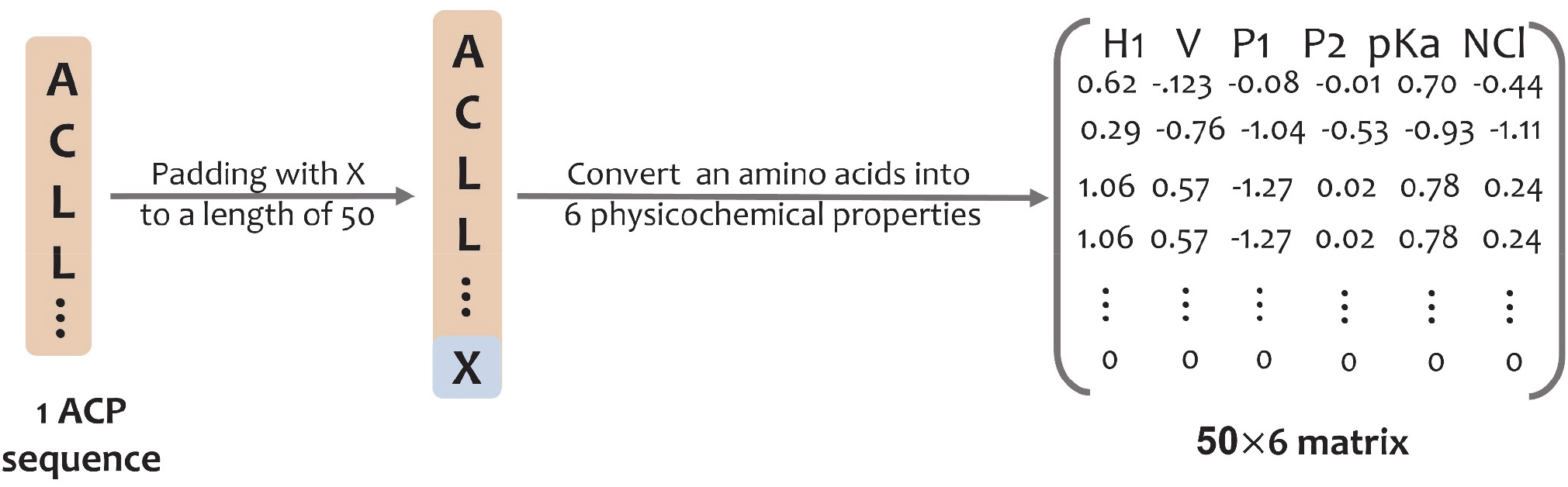
PC6 protein-encoding method. A padded ACP is transformed into a 50 × 6 matrix.

Subsequently, we implemented the neural network using Keras (https://github.com/keras-team/keras) from Tensorflow2 (https://www.tensorflow.org/). The model architecture consists of three blocks composed of convolutional layers, batch normalization, max pooling, dropout layers, and two dense layers (**Fig. 5**). The first dense layer contains 128 units with a 50% dropout rate. The last layer in the model is the output layer and is composed of a one-dimensional dense layer with the sigmoid activation function that produces a value ranging from 0 to 1; this value can indicate whether a peptide is an ACP. The convolutional layer in the three blocks in our model was built using 64, 32, and 8 one-dimensional filters of length 20 with the ReLU activation function, respectively. After the convolutional layer was built, batch normalization and max-pooling were applied with a 25% dropout rate in every block. Binary cross entropy was implemented as the loss function. With a learning rate of 0.0001, the Adam optimizer was used as our optimizer. We trained the model using the training data set (90%) and evaluated its performance using the validation data set (10%). Finally, all available data, namely 2124 positive and 2124 negative data, were used to train the final model.

**Fig. 5.**
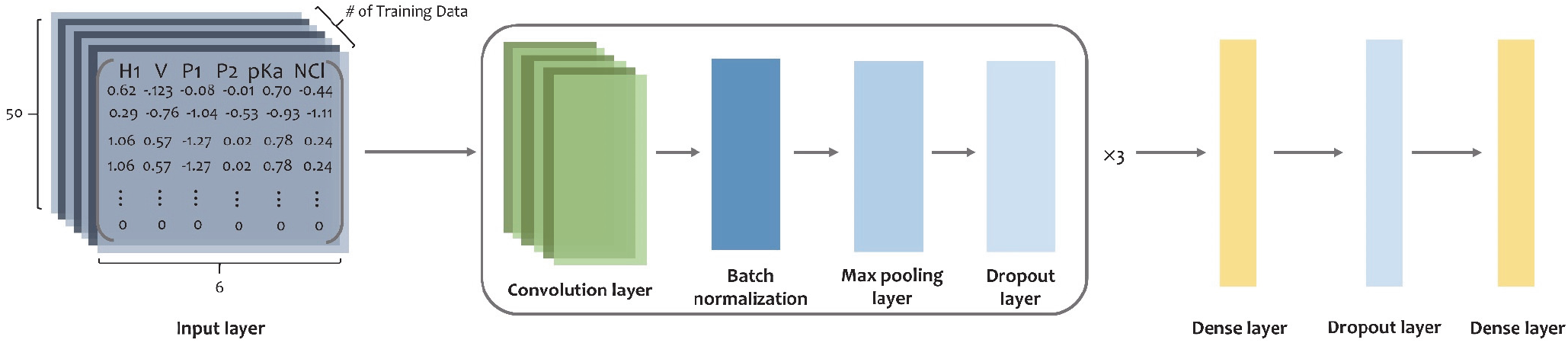
Model architecture in this study. After PC6 encoding, protein sequences go through every layer in this model.

### 1.4 Data for the final model

After we confirmed the most favorable model architecture and hyperparameters, we trained the model using all the available data (2124 positive and 2124 negative data) and eventually produced the final prediction model for the website. The data set used in this study can be found on our online HELP page. (https://axp.iis.sinica.edu.tw/AI4ACP/helppage.html) The positive and negative data sets will be continuously updated with the same criteria if new ACPs are discovered in the future.

### 1.5 System Implementation and Workflow

For the intuitive user experience and easy understanding, we built AI4ACP composed of the LAMP system architecture (Linux Ubuntu 16.04, Apache 2.04, MySQL 5.7, and PHP 5.1) with the Bootstrap 3 CSS framework (http://getbootstrap.com/), jQuery1.11.1, and jQuery Validation version 1.17. Furthermore, the core of the analysis process was implemented in the neural network by using Keras from Tensorflow. AI4ACP runs as a virtual machine (CPU of 2.27 GHz, 20 cores, 32-GB RAM, and 500-GB storage) on the cloud infrastructure of the Institute of Information Science, Academia Sinica, Taiwan.

AI4ACP is a website service that allows users to predict whether a query peptide sequence is an ACP. The input data should be in the FASTA format, and the query peptide sequence should be composed of only 20 essential amino acids; sequences would not be recognized if they contain unusual amino acids such as B, Z, U, X, J, or O. AI4ACP would output a CSV file containing a prediction score ranging from 0 to 1 and the prediction result as YES or NO for each input peptide sequence. The prediction score represents the probability that the query peptide sequence is an ACP. The prediction results shown as a binary column in the output file indicate the ACP sequence(s). The prediction result is based on the prediction score with a threshold of 0.472, which is the average of thresholds calculated by training the model five times. The workflow of AI4ACP is presented in Fig. 6 and explained as follows: First, the query peptide sequence is input in the FASTA format or as a FASTA file, and a valid job title is provided (Fig. 6A). After the query sequence is submitted, the result appears in a three-column table composed of the input peptide’s name, prediction score, and prediction result (Fig. 6B). In addition, a piechart presents the prediction result; this pie chart enables users to view the prediction results of the whole submission at the same time (Fig. 6C).

**Fig. 6.**
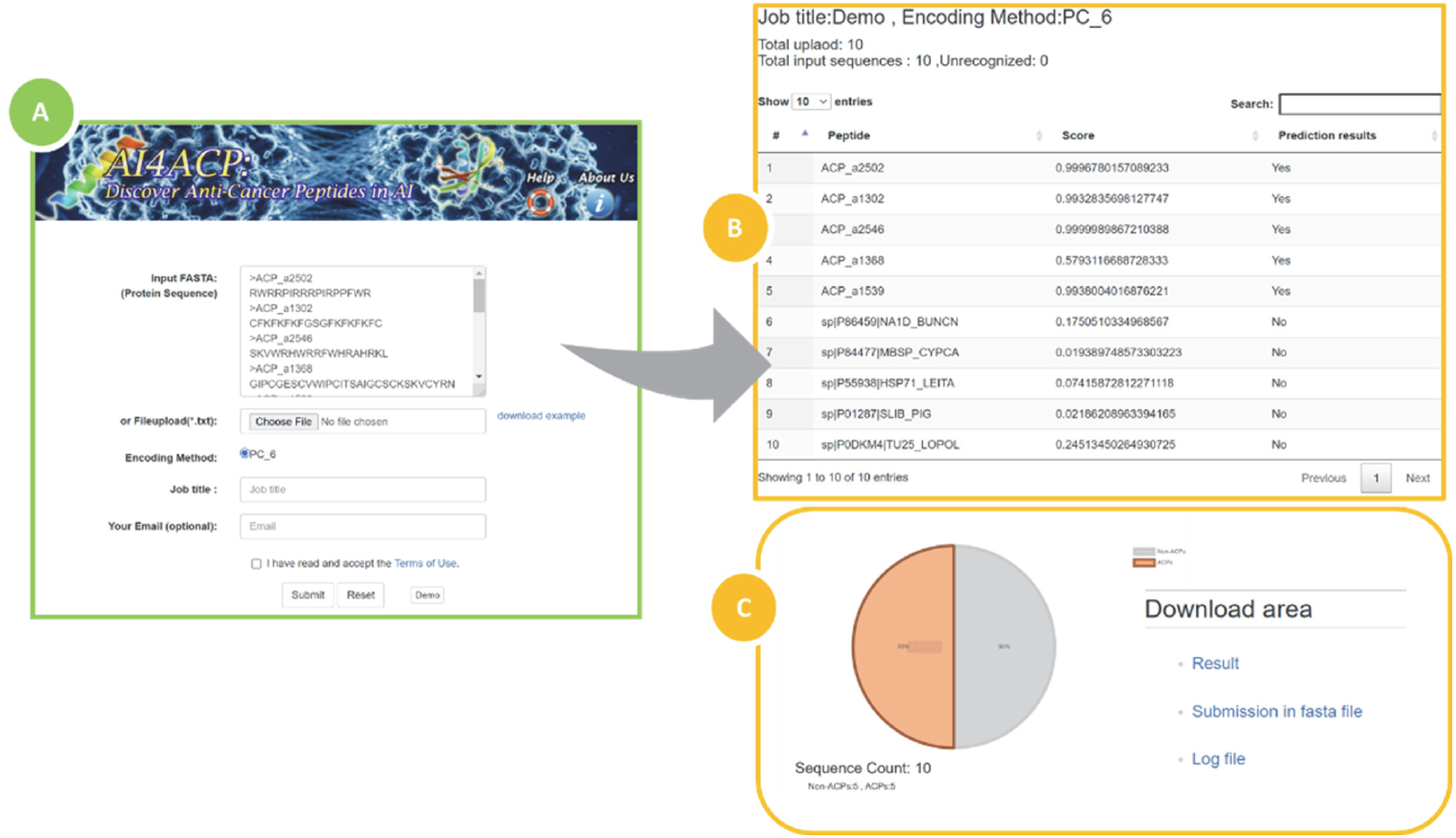
AI4ACP website. **A)** Web portal of AI4ACP for sequence submission in FASTA. **B)** Output of ACP activity for each submitted sequence with a prediction score. **C)** A piechart presenting the prediction of the whole submission. Moreover, the submission with files generated during the prediction.

## Results

We compared the performance of our model with those of other state-of-the-art ACP predictors. A previous study [14] indicated that most of the ACP predictors were trained and tested using two data sets: the main data set and an alternative data set. The main data set was 861 ACPs as the positive set and 861 AMPs as the negative set (80% for training and 20% for testing). The alternative data set consisted of 970 ACPs as the positive set and 970 peptide sequences randomly chosen from Swiss-Prot as the negative set (80% for training and 20% for testing).

In addition to these two data sets, we obtained a new collection. Table 1 shows the comparison of the composition of the data sets, and Fig. 7 shows the Venn diagram of the positive set of the data sets.

**Table 1.** Comparison of the composition of three data sets.

| Dataset |  | Positive set | Negative set |
| --- | --- | --- | --- |
| **Main set**<br>$(P^{861} + N^{861})$ | Training set | 689 ACPs | 689 AMPs |
|  | Testing set | 172 ACPs | 172 AMPs |
| **Alternative set**<br>$(P^{970} + N^{970})$ | Training set | 776 ACPs | 776 peptides from Swiss-Prot |
|  | Testing set | 194 ACPs | 194 peptides from Swiss-Prot |
| **New collection**<br>$(P^{2124} + N^{2124})$ | Training set | 1912 ACPs | 956 peptides from UniProt + 956 randomly generated sequences |
|  | Testing set | 212 ACPs | 106 peptides from UniProt + 106 randomly generated sequences |

**Fig. 7.**
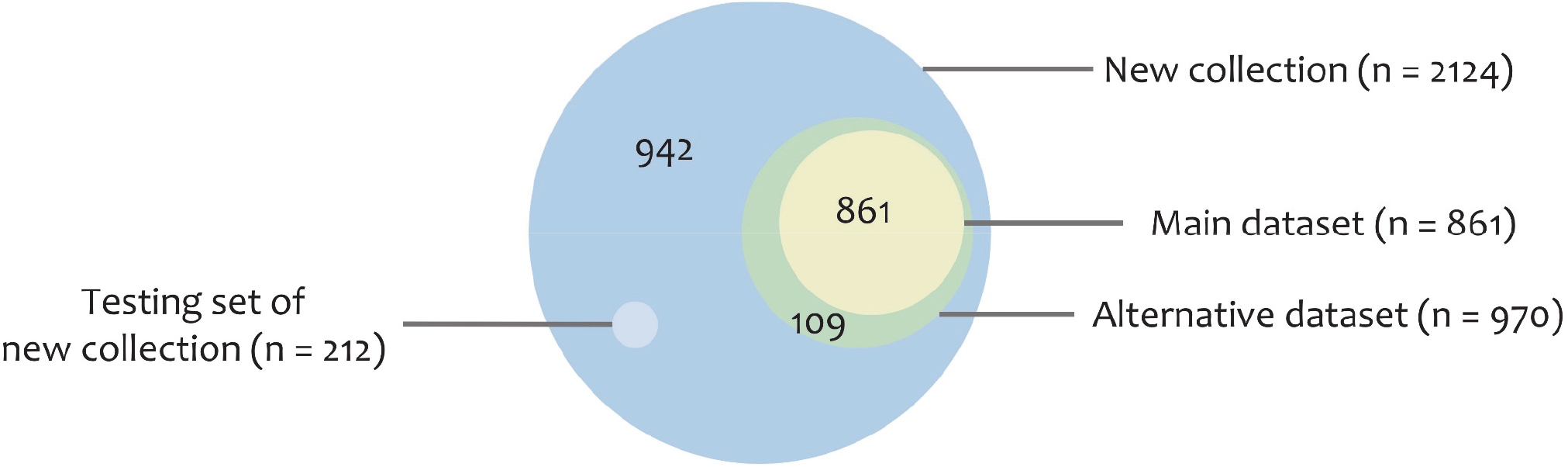
Venn diagram of the positive data sets.

We compared AI4ACP, trained using the main data set and the alternative data set in the previous study, with other state-of-the-art ACP predictors. Most of the ACP predictors lack maintenance, and thus they were not working. The results shown in Tables 2 and 3 were obtained from the published findings of AntiCP2.0 [16] and ACPred [14]. Table 2 presents the performance of ACP predictors trained and tested using the main data set. Most of the ACP predictors trained with the main data set did not perform efficiently. We observed that the specificities of most of the ACP predictors were not as satisfactory as their sensitivities. The composition of the negative main data set might have affected the ACP prediction performance of the ACP predictors. Because ACPs are a subset of AMPs, some of the AMPs may possess anticancer activity; thus, using AMPs as the negative set would be inappropriate. Thus, the ACP predictor was trained and tested in this study using the alternative data set.

**Table 2.**
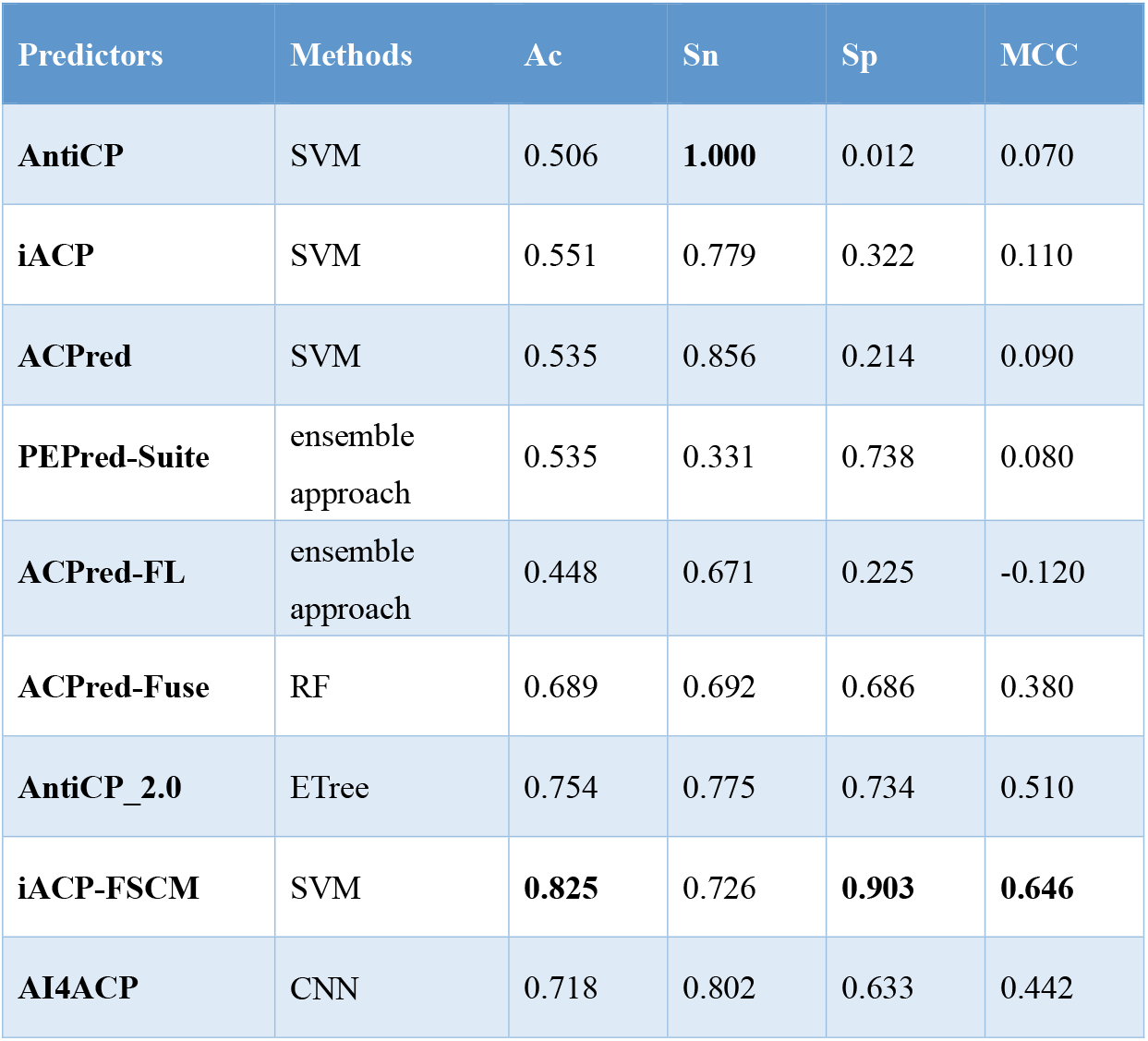
Comparison of ACP predictors trained and tested with the **main data set**. Results were obtained from the published findings of AntiCP2.0 and ACPred.

| Predictors | Methods | Ac | Sn | Sp | MCC |
| --- | --- | --- | --- | --- | --- |
| **AntiCP** | SVM | 0.506 | **1.000** | 0.012 | 0.070 |
| **iACP** | SVM | 0.551 | 0.779 | 0.322 | 0.110 |
| **ACPred** | SVM | 0.535 | 0.856 | 0.214 | 0.090 |
| **PEPred-Suite** | ensemble approach | 0.535 | 0.331 | 0.738 | 0.080 |
| **ACPred-FL** | ensemble approach | 0.448 | 0.671 | 0.225 | -0.120 |
| **ACPred-Fuse** | RF | 0.689 | 0.692 | 0.686 | 0.380 |
| **AntiCP_2.0** | ETree | 0.754 | 0.775 | 0.734 | 0.510 |
| **iACP-FSCM** | SVM | **0.825** | 0.726 | **0.903** | **0.646** |
| **AI4ACP** | CNN | 0.718 | 0.802 | 0.633 | 0.442 |

**Table 3.** Comparison of ACP predictors trained and tested using the **alternative data set**. Results were obtained from the published findings of AntiCP2.0 and ACPred.

| Predictors | Classifier | Ac | Sn | Sp | MCC |
| --- | --- | --- | --- | --- | --- |
| **AntiCP** | SVM | 0.900 | 0.897 | 0.902 | 0.800 |
| **iACP** | SVM | 0.776 | 0.784 | 0.768 | 0.550 |
| **ACPred** | SVM | 0.853 | 0.871 | 0.835 | 0.710 |
| **PEPred-Suite** | ensemble approach | 0.575 | 0.402 | 0.747 | 0.160 |
| **ACPred-FL** | ensemble approach | 0.438 | 0.602 | 0.256 | -0.150 |
| **ACPred-Fuse** | RF | 0.789 | 0.644 | **0.933** | 0.600 |
| **AntiCP_2.0** | ETree | **0.920** | **0.923** | 0.918 | **0.840** |
| **iACP-FSCM** | SVM | 0.889 | 0.876 | 0.902 | 0.779 |
| **AI4ACP** | CNN | 0.894 | 0.871 | 0.918 | 0.790 |

**Table 3** shows the performance of ACP predictors trained and tested using the alternative data set. The performance of most of the ACP predictors was more favorable than those trained using the main data set; AntiCP2.0 [16] exhibited the highest performance. In this study, AI4ACP was constructed using a deep learning model and thus required more data to improve the prediction accuracy.

Most of the state-of-the-art predictors lacked maintenance and were thus unable to predict the testing set of the new collection. AI4ACP and AntiCP2.0 [16], the only state-of-the-art web-based ACP predictors available, were evaluated using the testing set of the new collection. AI4ACP was trained using the alternative data set and the new collection with the model as mentioned earlier architecture, respectively, and both were tested using the testing set of the new collection. As shown in Table 4, the performance of our model was similar to AntiCP2.0 when trained using the alternative set but more favorable than that of AntiCP2.0 when trained using the new collection data set.

**Table 4.** Comparison of ACP predictors tested using the testing set of the new collection.

| Predictors | Classifier | Training set | Ac | Sp | Sn | MCC |
| --- | --- | --- | --- | --- | --- | --- |
| **AntiCP2.0** | ETree | Alternative set | 0.792 | 0.717 | **0.868** | 0.592 |
| **AI4ACP** | CNN | Alternative set | **0.802** | **0.750** | 0.854 | **0.607** |
| **AI4ACP** | CNN | New collection | **0.913** | **0.925** | **0.901** | **0.826** |

## Conclusion

The identification and screening of novel ACPs in a wet lab is usually time-consuming and expensive. Exploring the anticancer activity of peptides by using ACP predictors can accelerate the development of new anticancer drugs. However, the prediction of an ACP predictor is merely speculative. Laboratory experiments would still be required to confirm whether a peptide sequence possesses anticancer activity.

The results revealed that combining the PC6 encoding method and deep learning model could efficiently predict ACPs. The PC6 encoding method could exactly preserve the physicochemical properties of amino acids from original peptide sequences, and the deep learning model could learn these preserved features. In addition, with an increase in the number of peptide sequences confirmed as ACPs, we could build a predictor that exhibited more favorable performance and higher accuracy than other state-of-the-art ACP predictors. AI4ACP is a user-friendly web-based ACP predictor, and users can use this tool to detect whether the query sequence is an ACP. This tool can be beneficial for drug development for cancer treatment. AI4ACP will be continuously updated once new ACPs are discovered in the future. Besides, the deep learning model is available at https://github.com/yysun0116/AI4ACP.

## Abbreviations

Ac: Accuracy
ACPs: Anticancer peptides
AMPs: Antimicrobial peptides
CNN: Convolution neural networks
DNN: Deep neural networks
LAMP: Linux Ubuntu 16.04, Apache 2.04, MySQL 5.7, PHP 5.1
MCC: Matthew’s correlation coefficient
ReLU: Rectified linear unit
RF: Random Forest
Sn: Sensitivity
Sp: Specificity
SVM: Support vector machine

## Declarations

### Ethics approval and consent to participate

Not applicable

### Consent for publication

Not applicable.

### Availability of data and material

AI4ACP (web-server) with the dataset used is freely accessible at https://axp.iis.sinica.edu.tw/AI4ACP/. Furthermore, the model with code is also available on Github at https://github.com/yysun0116/AI4ACP.

### Authors’ contributions

YYS, TTL, and CYL collected the data, planned and implemented the algorithm. YYS, IHL, and CYL, composed the whole infrastructure, conducted the experiments, and drafted the manuscript together with TTL and CYL. WCC, IHLand SHC worked on constructing workflow and web platforms for data visualization and analysis. All of the authors had read and approved the final manuscript.

## Acknowledgments

Wallace Academic Editing edited this manuscript.

